# Optical crosstalk in SPAD arrays for high-throughput single-molecule fluorescence spectroscopy

**DOI:** 10.1101/207118

**Authors:** Antonino Ingargiola, Maya Segal, Angelo Gulinatti, Ivan Rech, Ivan Labanca, Piera Maccagnani, Massimo Ghioni, Shimon Weiss, Xavier Michalet

## Abstract

Single-molecule fluorescence spectroscopy (SMFS), based on the detection of individual molecules freely diffusing through the excitation spot of a confocal microscope, has allowed unprecedented insights into biological processes at the molecular level, but suffers from limited throughput. We have recently introduced a multispot version of SMFS, which allows achieving high-throughput SMFS by virtue of parallelization, and relies on custom silicon single-photon avalanche diode (SPAD) detector arrays. Here, we examine the premise of this parallelization approach, which is that data acquired from different spots is uncorrelated. In particular, we measure the optical crosstalk characteristics of the two 48-SPAD arrays used in our recent SMFS studies, and demonstrate that it is negligible (crosstalk probability ≤ 1.1 10^−3^) and is undetectable in cross-correlation analysis of actual single-molecule fluorescence data.

## 1. Introduction

Single-molecule fluorescence spectroscopy includes several powerful techniques used to investigate fundamental physical, chemical and biological processes at the molecular level [2, 3, 4]. In its freely-diffusing molecule version, where molecules briefly transit (in a few ms) in and out of a confocal excitation spot, concentration needs to be kept low so that individual molecule “bursts” of photons can be identified. This results in long acquisition time (10 min to several hr) in order to accumulate sufficient statistics. To overcome this limitation, we use a multispot excitation/detection approach, and matching arrays of silicon single-photon avalanche diodes (SPAD) arrays [5, 6, 7, 1]. The number of photons in a burst is limited by the diffusion speed of typical biomolecules across the excitation volume, by the excitation power and by the photophysical properties (e.g. brightness) of the fluorophores used to label molecules of interest. Because of the low photon-count (∼ 100) of typical bursts, SMFS requires SPADs with high photon-detection efficiency (PDE) in the visible range of the spectrum. For these reasons, our SPAD arrays are fabricated using a custom process, instead of the more common complementary metal-oxide semiconductor (CMOS) process [8, 9]. We recently demonstrated a SMFS system achieving close to 2-orders of magnitude throughput enhancement for single-molecule fluorescence resonant energy transfer (smFRET), using two excitation lasers and two 48-pixels SPAD arrays, to collect photons emitted by the “donor” and “acceptor” fluorophores attached to each molecule [1]. This approach can be used for real-time studies of kinetic processes and opens the way to applications of SMFS in medical diagnostics and drug discovery.

One of the assumptions of parallel data acquisition is that data acquired in each spot is uncorrelated (independent) from that acquired in others. In other words, each spot samples an independent volume of the solution of single molecules, and no molecules are observed in separate spots at different times. This assumption is verified in our experiments, where spots are separated by a few μm, which translates in negligible probability of observing a molecule in a nearby spot after it has been detected in an initial one. Note that molecules may get in and out of each individual spot a few times before wandering away, a recurrence phenomenon which can be used to study long term dynamics in single-molecules [10, 11].

A more subtle requirement for the independence of singlemolecule measurements performed in nearby spots, is the absence of crosstalk. We can define crosstalk between two spots as the presence of a fraction of the signal of spot A in the signal measured in spot B. One source of crosstalk between spots is due to the collection of light emitted from spot A by SPAD B, supposed to detect signal only from B. This can be readily excluded for similar geometrical reasons as used above to exclude long term correlation between spots. A less obvious but well-known type of cause (see section 2) specific to SPAD arrays, is the occurrence of internal electronic or optical crosstalk between the pixel pairs. Electrical crosstalk is usually negligible in a well designed SPAD array. Crosstalk between spots, regardless of its origin, does show up in cross-correlation functions, and can potentially affect some single-molecule analysis.

In this paper, we describe an accurate method for optical crosstalk estimation (section 2) and use it to characterize the optical crosstalk in the two SPAD arrays used in our 48-spot SMFS setup [1] (section 3). We show that the low crosstalk of these devices has a negligible effect on single-molecule measurements.

### Data and code availability

All raw data presented are freely available on Figshare (doi.org/10.6084/m9.figshare.5146096). Notebooks and code used for analysis and simulations are open source and available on GitHub github.com/tritemio/48-pixel-SPAD-crosstalk-analysis.

## 2. Optical crosstalk

Optical crosstalk between two SPADs within the same array can happen when an avalanche is triggered in one SPAD: hot carriers emit secondary photons [12, 13] which can propagate within the device structure, directly or after multiple refflections, and trigger an avalanche in another SPAD [14]. This secondary photons emission is proportional to the avalanche current [13] which typically lasts < 20 ns in SPADs using active quenching circuits [15]. Hence, when a crosstalk event occurs, the second SPAD is triggered with a delay < 20 ns. Optical crosstalk is a phenomenon present both in silicon (Si) custom [16], Si CMOS [17], III-V [18, 19] SPAD arrays and, particularly troublesome in Si photomultipliers (SiPM) [20, 21]. When the crosstalk probability is high (≥ 10%), cascades of multiple crosstalk events can be detected, although this is not usually an issue in Si SPAD arrays. In SiPM, where the signal from individual SPADs cannot be measured independently and the large crosstalk probability makes cascade effects nonnegligible, optical crosstalk characterization is complex and requires sophisticated models [22, 21]. By contrast, in SPAD arrays, the crosstalk probability can be easily estimated by simple coincidence measurements as described next.

The crosstalk probability *P_c_* can be estimated from the number of “coincidence events”, that is the number *C* of photons recorded within a given (short) time window Δ*t* by two pixels *A* and *B*, over an acquisition duration *T*. Usually, crosstalk is estimated from dark counts acquisition, although, to reduce the acquisition time, SPADs can be also illuminated with a constant signal. The number of coincidences C can be measured with a dedicated asynchronous circuit [16, 17] or using the photon timestamps obtained with a simple counting board as used in single-molecule measurements. In this case, Δ*t* needs to be chosen larger than the avalanche duration plus the time-stamp resolution (typically 10-20 ns).

The number of coincident events due to uncorrelated counts *C_u_* can be computed from Poisson statistics, which we will assume to describe dark counts (similar expressions can be obtained for other statistics). Given two Poisson random variables *X*_A_ ∼ Poi{λ_A_Δ*t*} and *X*_B_ ∼ Poi{λ_B_Δ*t*} with mean event rate per unit time λ_A_ and λ_B_, the probability of having at least one count in each variable during a time window Δ*t* is:

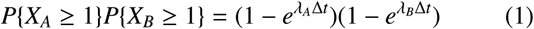

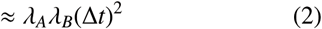

where the approximation holds when Δ*t* ≪ λ^−1^. Hence, for two uncorrelated Poisson processes, the number of coincidences in Δ*t* during a time *T* is:

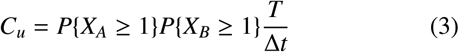

The number of crosstalk events *C_c_* can be computed as:

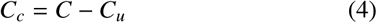

and the crosstalk probability as:

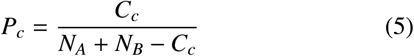

The standard deviation of the *P_c_* estimator in eq. 5 can be computed from the standard deviation of *C_c_*, which is a Poisson random variable as well (*C_c_* counts are independent events):

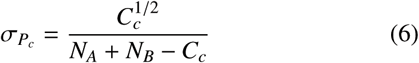

The rates λ_A_ and λ_B_ in eq. 1 can be estimated from dark counts measurements as:

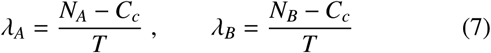

where *N_A_* and *N_B_* are the total counts in SPAD A and B during duration *T*. Using eq. 7 in eq. 4 leads to an equation that can be solved for *C_c_*, so that the probability of crosstalk and its standard deviation can be computed from eq. 5 and 6. The supporting notebook *Crosstalk Theory and Simulations* reports crosstalk simulations confirming that estimator in eq. 5 is unbiased.

## 3. Results and Discussion

We used eq. 1, 5, 6 and 7 with Δ*t* = 50 ns to compute the crosstalk in the two 48-SPAD arrays used in ref. [1]. These arrays have a 12 × 4 geometry with 500 μm pitch and 50 m diameter. In order to achieve accurate estimation, even for SPADs with very low dark count rates (DCR) (∼50 cps, see ref. [1] for details), we acquired data for ∼6 hours. Computational details can be found in the supporting notebook *Optical crosstalk estimation*.

Results in Fig. 1 show that the optical crosstalk probability *P_c_* is ∼ 1.1 10^−3^ for nearest-neighbor SPADs, ∼ 1.5 10^−4^ for nearest-neighbor SPADs on a diagonal, and decreases by more than one order of magnitude down to negligible levels for SPADs farther apart. The crosstalk dependence with distance (Fig. 1, Panel A & D) indicates a much stronger absorption of secondary photons within the device compared to previous arrays [7, 14]. An *R*^−2^ dependence (black dashed curve) is indicated for comparison and corresponds to a simple isotropic attenuation model, assuming direct propagation between pixels and no absorption. By contrast, crosstalk was observed to decrease slower than this simple *R*^−2^ model in a previous generation of SPAD arrays, an effect attributed to reflection of near infrared photons of the chip’s bottom surface [14]. The stronger attenuation observed here has been obtained by fabricating the arrays on a silicon substrate with an especially high doping level (> 2 10^19^ cm^−3^). At these concentrations, free carriers absorption strongly reduces the propagation of near infrared photons through the substrate [23]. The good uniformity of crosstalk probability across both arrays also suggests that no edge effect is present in these devices, in contrast with the previous generation of devices [14]. Reflections of the chip edges, whose minimum distance from a SPAD is 450 μm, are made negligible by the strong reduction of the crosstalk with the distance.

**Figure 1:**
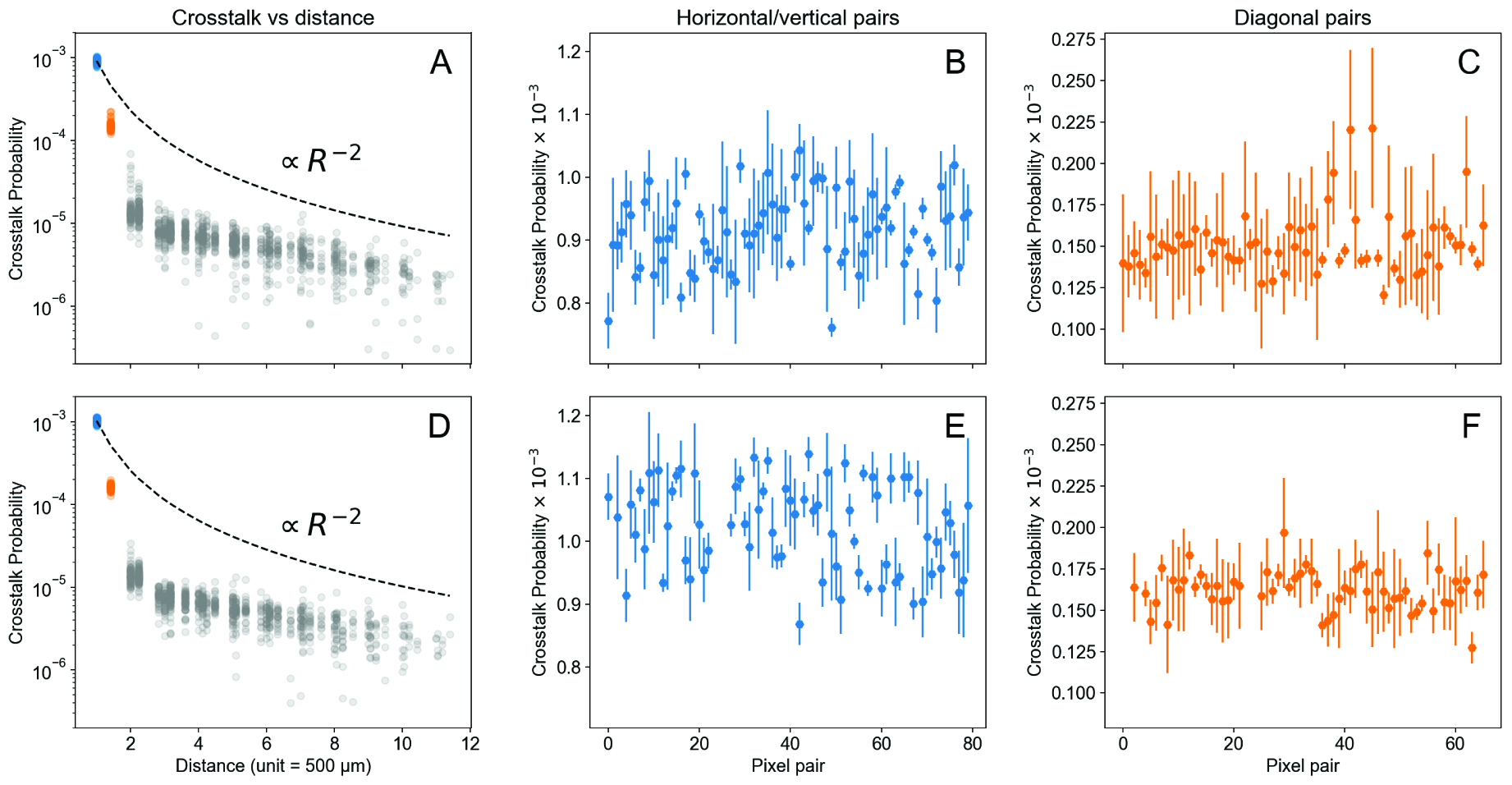
Optical crosstalk probability in the two 48-SPAD arrays used in our experiments. The two rows (A-C) and (D-F) present data from the “acceptor” and “donor” SPAD array respectively. Panels A,D show the crosstalk probability as a function of the distance between all pixel pairs. The black dashed line is a curve proportional to *R*^−2^ passing through the mean crosstalk of the nearest neighbor pairs (*blue dots*). The first two distances are color-coded and reported in detail in Panels B,E (nearest-neighbor pairs, 500 μmm: *blue*) and Panel C,F (nearest diagonal neighbor pairs, 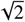 500 μm: *orange*). Error bars in Panels B,C,E,F indicate the ±3σ range, with σ computed according to eq. 6.

These excellent crosstalk characteristics can be further observed in Fig. 2, which reports the auto-correlation and crosscorrelation functions (ACF & CCF, eq. 8) of the donor intensity in 4 adjacent pixels during a typical smFRET measurement [1]. The curves have been computed with the open source Python package pycorrelate, using the algorithm from ref. [24]. See the supporting notebook *ACF and CCF* for details.

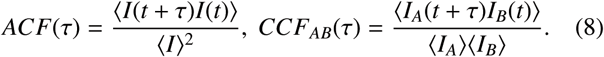

Assuming an ideal Gaussian cylindrical profile for the excitation/detection probability, the ACF of a solution of freely diffusing molecules at equilibrium takes a simple form given by eq. 9 [25, 26].

**Figure 2:**
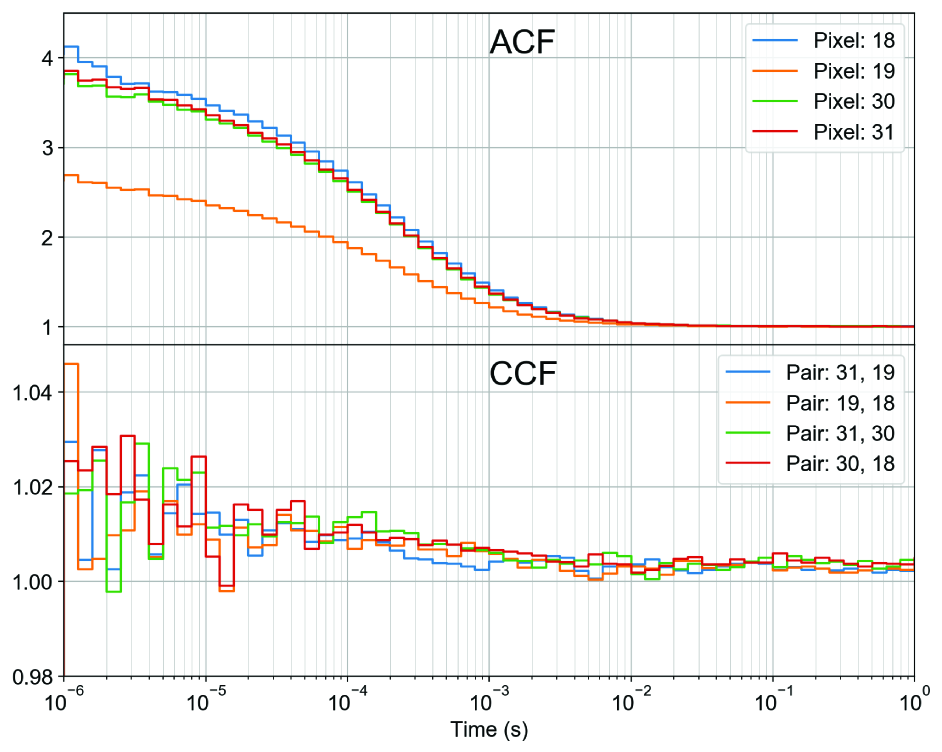
Normalized intensity auto- and cross-correlation functions for a group of 4 adjacent SPADs during a typical smFRET experiment.

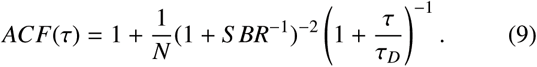

In eq. 9, *N* is the average number of molecules present in the excitation/detection volume and *S BR* is signal-to-background ratio. This function exhibits a characteristic inflexion point around the diffusion time through the excitation/detection volume, *τ_D_*, and ideally plateaus to a value 1/*N* at short time scale. In practice, additional contribution of the SPAD afterpulsing at short time scale (not shown) mask this asymptotic behavior [5].

In the absence of measurable molecular transport from one spot to another, no correlation between the signals measured in the two different spots A and B is expected, and the CCF of the intensities recorded in the two spots, *CCF _AB_* (τ) (eq. 8) should be equal to 1 (at least at time scales larger than ∼ 1 μs). In the presence of crosstalk, an “echo” of the ACF of each spot is expected, whose amplitude is that of the ACF (1/*N* in eq. 9) multiplied by the crosstalk probability, *P_c_*. Fig. 2 is compatible with this prediction, the CCF amplitude (*P_c_*/*N*) being smaller than the statistical noise level present in these (relatively) short measurements.

## 4. Conclusions

In this study, we have characterized the optical crosstalk of two large SPAD arrays used in single-molecule fluorescence experiments, and showed it to be extremely low. In particular, we have verified that it does not affect CCF analysis, a particularly useful property for high-throughput measurements in flow geometries. The ability to perform CCF analysis between distinct spots with negligible influence of crosstalk makes it practical to use these SPAD arrays for high-throughput SMFS measurements in flow geometries, where transport of molecules from one spot to another, and conformational change during that time, would result in non-trivial CCF signatures. Future work will take advantage of this possibility.

## Acknowledgments

This work was funded by NIH Grant R01 GM095904. S. Weiss discloses intellectual property used in the research reported here. The work at UCLA was conducted in Dr. Weiss’s Laboratory. M. Ghioni discloses equity in Micro Photon Devices S.r.l. (MPD). No resources or personnel from MPD were involved in this work. We are grateful to our colleagues at UCLA and Polimi for their contributions to this project over the years.

